# The perceptions and roles of teachers and caregivers on play in promoting children’s early learning and development in a rural Kenya

**DOI:** 10.1101/2024.12.17.629004

**Authors:** Paul Otwate, Margaret Nampijja, Nelson Langat, Linda Oloo, Silas Onyango, Jesse Mabongah, Erick Makhapila, Silas Ooko, Brian Odhiambo, Patricia Kitsao-Wekulo

## Abstract

Adults, including teachers and parents have an important role in supporting children’s learning and development through play. This paper aimed to explore parents’ and teachers’ viewpoints and roles on play in promoting early learning and development in children, which has little emphasis in existing studies. This paper employed a cross-sectional study using a mixed-method approach to gather data from preschool teachers (n=96), parents (n=126), and policy implementers (n=6) in Kenya. We used questionnaires, focus group discussions, and interview guides to collect qualitative and quantitative data through face-to-face interviews with teachers and parents and mobile interviews with policy stakeholders. We performed qualitative analysis through reviewing the study transcripts, coding, and generating emerging themes that later guided the draft of the narrative report. Quantitative data were coded, cleaned, and reviewed before they were descriptively computed on the Stata version 18 to generate graphic representations. Our findings revealed that teachers and parents jointly perceived play to be pivotal on children’s learning and development. In addition, parents and teachers recognized that it is their role to engage children in play, provide play materials, and supervise children. Based on our results, we suggest that preschool teachers and parents collaboratively identify days to jointly develop play materials in preschool centers. This strategy may present a better understanding of the adults’ primary role of facilitating play and play-based learning. Our findings may serve as a resource and source of information for preschool teachers, parents, and policy stakeholders to consider play not only as a development pathway but also an element of learning and contributor to school readiness in children.

## Introduction

Play has a vital role social, physical, language, and cognitive development in children. Despite the highly contested conceptualizations about play, play-based learning and pedagogy in early childhood care and education, play is crucial in contributing to learning, development, and school readiness in children (McLean et al., 2022; Zosh, Hopkins, Jensen, Liu, Neale, Hirsh-Pasek, et al., 2017). There is evidence that play-based learning supports learners in development of communication, collaboration, cognitive, social, emotional, and fine motor skills (Parker et al., 2022). Notwithstanding the factors contributing to the decisions adults make on provision of play, the learner-centered approaches of play characterized by joyful, meaningful, socially engaging, and interactive have far reaching impact on school readiness in children (Parker et al., 2022; Zosh, Hopkins, Jensen, Liu, Neale, Hirsh-pasek, et al., 2017). Teachers, parents, and relevant caregivers are critical in providing a quality play-based learning environment for learners to acquire the needed competencies that align with the sustainable development goal (SDG) of education number 4 (Tadesse et al., 2023). Correct conceptualization of play and play-based learning will influence the decision significant adults make about provision of play in early childhood education.

Although play is considered an essential strategy in promoting preschool and school readiness, understanding the significant roles and perceptions of parents and teachers may lead to its fruition (Tadesse et al., 2023). Transforming the perceptions of teachers and parents to take leading roles in promoting play-based experiences becomes crucial in contemporary research (Aronsson, 2009). In a study done in Palestine by Murtagh (2022), caregivers needed adequate knowledge to conceptualize play as a meaningful aspect linked to children’s learning and development outcomes (Khalil et al., 2022). Another study in the same country attributed the integration of learning through play approaches to improved academic achievement in Mathematics activities in learners (Murtagh et al., 2022; Tadesse et al., 2023). For instance, learners in the intervention group scored higher (P < 0.01) due to the support teachers had provided on meaningful use of play-based learning approaches. This finding suggests a positive relationship between parents’ attitudes and perceptions and that of teachers in promoting use of play-based approaches (Gracekabura, 2020).

Even though play-based learning contributes to holistic development in children, cultural factors may shift the “traditional” or more “highly guided” teacher-centered approaches commonly used at the primary school level (Taiwo Ogunyemi & Henning, 2020; Thomsen & Stjerne, 2019). For instance, while using a different approach of Quasi-experimental evaluation in Bangladesh through LtP, children who scored low at baseline were able to catch up to their peers who initially had higher scores, showed increased knowledge and skills, and demonstrated improved quality of their interactions with others as a result of perceived benefits of play-based learning in teachers (Thomsen & Stjerne, 2019). Similarly, a randomized control trial (RCT) study in rural Ghana found that LtP improved learner’s emerging literacy, executive functioning, and fine motor skills largely due to parents’ positive perceptions of the role of play (Rother et al., 2020). Positive perceptions about play may influence the way teachers and parents view play and degree to which they would value playtime.

Study findings by Ngware et al. (2018) and Ndlovu et al. (2023) revealed negative perceptions by school practitioners in rural areas who in particular had insufficient understanding of the use of play as an instructional pedagogy in early-grade learners (Ngware et al., (2018); Ndlovu et al., 2023). These findings are relevant to this paper because they demonstrate an urgent need to empower teachers and improve their perceptions about play-based learning in early childhood education. Consequently, young children learn best when caregivers demonstrate a positive perception of the play and playful instructional approaches within their learning environment (UNESCO, 2022). In Kenya, play-based approaches are explicit in the implementation of competency based curriculum (CBC) to promote main core learning areas including problem-solving and critical thinking skills (Syomwene & Mwaka, 2018).

Therefore, giving an insight into the significance of positive perceptions in parents, caregivers, and teachers can enhance a holistic viewpoint of play as an approach that not only promotes physical skills but also enhances health, social, cognitive, language, and emotional milestones in young children (Ducusin & Dy, 2016). The success in these milestones may certainly become requisite for a smooth transition from home to preschool and later into primary level (Ducusin & Dy, 2016). Limited studies in sub-Saharan Africa have underpinned a clear understanding of the attitudes and knowledge of significant adults into promoting play-based learning, with little evidence to demonstrate the influence of perceptions and primary roles of parents and those of teachers in preschools affect the decisions they make on provision of less structured play experiences (Rother et al., 2020; Syomwene & Mwaka, 2018; Tadesse et al., 2023).

In Kenya, parents, and teachers play crucial roles in contributing to playful learning experiences in young children at home and school. They are considered enablers of successful transition from home to preprimary schooling and subsequent grades (Syomwene & Mwaka, 2018). Whereas some educators consider LtP as an adjunctive strategy to be used in conjunction with more conventional, rote approaches to teaching and learning (Lenjebo et al., 2023), there is a dearth of evidence if the relationship between parents’ and caregivers’ perceptions play a role transforming this school of thought in sub-Saharan Africa. Moreover, although research have explored parents’ and teachers’ attitudes towards play, scant mixed-method research has focused to establish a linkage between the perceptions and roles of parents and teachers in contextualizing and conceptualizing the place of play and play-based learning in public preschools in SSA. Therefore, this paper seeks to add and explore new knowledge to explicitly understand the influence of the perceptions and roles of parents and teachers in detail on play and play-based learning as a predictor of children’s early learning, development, and school readiness. This study purposively aimed to; determine teachers’ perceptions of the benefits of play, parents’ perceptions of the benefits of play in promoting children’s learning and development, and to examine the significant roles of parents and teachers in the implementation of play-based learning approaches in public and private preschools in Kenya.

## Materials and Methods

### Study design

This paper employs a cross-sectional study adopting a mixed–method approach combining quantitative and qualitative techniques.

### Study context and site

This research was part of a larger study namely strengthening the capacity of teachers on play-based learning: evidence from Kenya, which was conducted in three counties, namely Kajiado, Kiambu, and Nairobi Counties. Kajiado County, with its nomadic pastoralist community and low levels of education, provided a unique rural context for exploring the perceptions of parents and teachers on play and play-based learning. Kiambu County was chosen for its peri-urban setting, characterized by a diverse population living on the outskirts of urban centers and rural areas. Nairobi City County was selected to represent an urban setting, with the study focusing on Makadara sub-County, which comprises Kenya’s largest informal settlements and middle-income residential areas. These mixt contexts presented opportunities for this study to explore the link between teacher’ and parents roles and perception in promoting play-based learning.

### Study participants

Participants were recruited from public and private preschools (n=30) in the three counties, 10 from each county comprising 96 preschool teachers and 126 parents. The recruitment of participants began on 23^rd^ November 2022 and ended on December 12, 2024 for the baseline and endline data collection respectively. We also interviewed two Early Childhood Development and Education (ECDE) officials from each of the three counties. 100 participants in the qualitative component were selected using purposive sampling and later interviewed.

### Data collection procedures

This paper uses data from the Knowledge and Innovation Exchange (KIX) study which interviewed 126 parents and 96 teachers from 30 pre-schools in Kenya.

In preparation for the data collection, 18 qualified field interviewers were subjected to a one-week training. On the last day of the training, a pre-test was conducted to assess the reliability of the test items, and appropriateness of the questions to participants, gauge the interviewers’ competence in conducting the interviews, and correct any issues with the study tools.

Our study participants provided written consent for the entire data collection exercise comprising qualitative and quantitative data sets. We submitted our consent forms together with the study protocol for approval by the Amref Scientific Ethics and Review Committee for approval before commencing our study. Our study did not include minors. In this respective, field interviewers with college-level education and experience in qualitative interviews conducted the focus group discussions (FGDs). Key informant interviews (KIIs) were carried out by members of the research team. Additionally, a field supervisor with a relevant college-level education was appointed to oversee the fieldwork and ensure data quality through spot checks.

Quantitative data were captured electronically using tablets after which they were transmitted to the APHRC SurveyCTO server. Face-to-face interviews with teachers and parents were conducted at their respective schools. On average, the interviews took about 45 minutes to complete. The qualitative interviews took between one and half hours to complete. The interviews were audio recorded and transcribed verbatim. Endline data were collected between November and December 2023.

### Data quality and management

The data were uploaded to the APHRC server, accessible only by the research team. Trained team leaders, with oversight from the APHRC research team, supervised the data collection exercise. Field team leaders verified the completeness of the data before uploading them to the server. Discrepancies were rectified by the team leader and the APHRC team during the daily briefing after the data collection exercise. To enhance the integrity of the data, the research team conducted regular spot checks and sit-ins during the data collection exercise.

The quality of the qualitative data was enhanced by recruiting and training qualified field interviewers with experience in qualitative data collection. The research team reviewed samples of the qualitative transcripts to ensure all the items were included as reflected in the FGD and KII guides. The interviews were transcribed verbatim by an experienced transcriber.

### Data analysis

The extracted quantitative data were cleaned and analyzed using Stata v17. The descriptive statistics were computed and presented using tables to establish the distribution of demographic factors, while the roles and perceptions of teachers and parents were presented using tables and bar graphs.

### Ethical considerations

The Knowledge and Innovation Exchange (KIX) study, a genesis of this paper, was internally approved by the APHRC’s Scientific Review Committee (SRC). Ethics renewal was obtained from Amref Health Africa’s Ethics and Scientific Review Committee (ESRC). The National Commission for Science, Technology, and Innovation (NACOSTI) granted us a permit to conduct the study. Written information consents were obtained from all participants before the interviews were conducted. We informed all the study participants that their participation was voluntary and that they were free to withdraw at any point during the study without any negative consequences. Further, we assured the participants that their identities and data would be kept anonymous and confidential.

### Data availability statement

The authors declare published data on this work will be available without restrictions to the public for consumption.

## Results

### Participants’ characteristics

The study team interviewed 96 preschool teachers and 126 parents for quantitative data and 66 parents and 42 teachers for qualitative data components from the 30 preschools.

### Caregivers’/parents’ socio-demographic characteristics

**Table 1** presents information on the socio-demographic characteristics of parents interviewed by county. Most parents had a mean age of 37.0 and were female (78%). Over 40% of interviewees from Kajiado had no education. On average, there were four children per household in Kajiado County, nearly a double percentage in Nairobi and Kiambu Counties.

**Table 1.** Parents’ socio-demographic characteristics.

|  | Kajiado |  |  | Kiambu |  |  | Nairobi |  |  |
| --- | --- | --- | --- | --- | --- | --- | --- | --- | --- |
|  | Baseline<br>N=44 | Endline<br>N=45 | p-value | Baseline<br>N=36 | Endline<br>N=42 | p-value | Baseline<br>N=39 | Endline<br>N=39 | p-value |
| Age (years) |  |  |  |  |  |  |  |  |  |
| Mean (SD) | 38.7<br>(10.7) | 39.5<br>(12.4) | 0.76 | 36.5 (8.7) | 35.9<br>(10.2) | 0.79 | 35.7 (7.2) | 34.6 (5.6) | 0.49 |
| Gender |  |  | 0.87 |  |  | 0.15 |  |  | 0.41 |
| Female | 32 (73%) | 32 (71%) |  | 24 (67%) | 34 (81%) |  | 29 (74%) | 32 (82%) |  |
| Male | 12 (27%) | 13 (29%) |  | 12 (33%) | 8 (19%) |  | 10 (26%) | 7 (18%) |  |
| Marital status |  |  | 0.55 |  |  | 0.29 |  |  | 0.80 |
| Married | 34 (77%) | 34 (76%) |  | 31 (86%) | 31 (74%) |  | 31 (79%) | 32 (82%) |  |
| Never married | 1 (2%) | 4 (9%) |  | 2 (6%) | 7 (17%) |  | 4 (10%) | 2 (5%) |  |
| Widowed | 7 (16%) | 5 (11%) |  | 0 (%) | 0 (%) |  | 1 (3%) | 2 (5%) |  |
| Divorced/<br>Separated | 2 (5%) | 2 (4%) |  | 3 (8%) | 4 (10%) |  | 3 (8%) | 3 (8%) |  |
| Highest level of<br>formal education<br>completed |  |  | 0.82 |  |  | 0.038 |  |  | 0.26 |
| None | 20 (45%) | 21 (47%) |  |  |  |  | 1 (3%) | 1 (3%) |  |
| Primary | 14 (32%) | 11 (24%) |  | 18 (50%) | 9 (21%) |  | 11 (28%) | 7 (18%) |  |
| Secondary | 6 (14%) | 6 (13%) |  | 8 (22%) | 15 (36%) |  | 7 (18%) | 16 (41%) |  |
| College | 3 (7%) | 4 (9%) |  | 7 (19%) | 16 (38%) |  | 15 (38%) | 12 (31%) |  |
| University | 1 (2%) | 3 (7%) |  | 3 (8%) | 2 (5%) |  | 5 (13%) | 3 (8%) |  |
| Occupation |  |  | 0.008 |  |  | 0.59 |  |  | 0.005 |
| Not employed | 16 (36%) | 4 (9%) |  | 8 (22%) | 12 (29%) |  | 10 (26%) | 6 (15%) |  |
| Employed<br>(Salaried) | 4 (9%) | 4 (9%) |  | 7 (19%) | 6 (14%) |  | 7 (18%) | 5 (13%) |  |
| Business | 7 (16%) | 20 (44%) |  | 7 (19%) | 12 (29%) |  | 15 (38%) | 5 (13%) |  |
| Farming | 7 (16%) | 9 (20%) |  | 6 (17%) | 3 (7%) |  | 0 (0%) | 1 (3%) |  |
| Employed<br>(Waged/Casual) | 10 (23%) | 8 (18%) |  | 8 (22%) | 9 (21%) |  | 7 (18%) | 22 (56%) |  |
| Number of<br>children |  |  |  |  |  |  |  |  |  |
| Mean (SD) | 4.5 (1.9) | 4.6 (2.1) | 0.85 | 2.8 (1.4) | 2.8 (1.2) | 0.96 | 2.6 (1.1) | 2.5 (1.1) | 0.84 |

### Teachers’ socio-demographic characteristics

Information on teacher’s socio-demographic characteristics are presented in **Table 2**. The mean age of teachers was about 42.0. Most of the teachers (97%) were female. Over 90% of the teachers had attained college-level formal education and were employed with the County Governments. Over 50% had more than five years of teaching experience across the three counties respectively.

**Table 2.** Teachers’ socio-demographic characteristics.

|  | Baseline<br>N=24 | Endline<br>N=23 | Baseline<br>N=31 | Endline<br>N=31 | Baseline<br>N=37 | Endline<br>N=42 |
| --- | --- | --- | --- | --- | --- | --- |
| Teacher's age (years); Mean (SD) | 41.2 (6.3) | 41.8 (6.9) | 41.6 (6.9) | 42.1 (6.8) | 42.3 (8.0) | 42.9 (8.6) |
| Gender |  |  |  |  |  |  |
| Female | 21 (88%) | 20 (87%) | 31 (100%) | 30 (97%) | 37 (100%) | 42 (100%) |
| Male | 3 (13%) | 3 (13%) | 0 (0%) | 1 (3%) | 0 (0%) | 0 (0%) |
| Position in school |  |  |  |  |  |  |
| Center manager | 4 (17%) | 1 (4%) |  |  | 10 (27%) | 10 (24%) |
| Head of ECD class | 6 (25%) | 3 (13%) | 10 (32%) | 8 (26%) |  |  |
| ECD teacher | 14 (58%) | 18 (78%) | 16 (52%) | 21 (68%) | 27 (73%) | 32 (76%) |
| Assistant teacher | 0 (0%) | 1 (4%) | 5 (16%) | 2 (6%) | 0 (0%) | 0 (0%) |
| ECDE classes currently taught in the center |  |  |  |  |  |  |
| Playgroup (0-3 years) | 0 (0%) | 0 (0%) | 0 (0%) | 0 (0%) | 3 (8%) | 3 (7%) |
| PP1 | 12 (50%) | 10 (43%) | 16 (52%) | 16 (52%) | 15 (41%) | 12 (29%) |
| PP2 | 12 (50%) | 10 (43%) | 15 (48%) | 15 (48%) | 19 (51%) | 24 (57%) |
| Both PP1 & PP2 | 0 (0%) | 3 (13%) | 0 (0%) | 0 (0%) | 0 (0%) | 3 (7%) |
| Highest level of formal education completed |  |  |  |  |  |  |
| Primary | 0 (0%) | 1 (4%) | 0 (0%) | 0 (0%) |  |  |
| Secondary | 0 (0%) | 0 (0%) | 0 (0%) | 0 (0%) | 1 (3%) | 4 (10%) |
| College | 23 (96%) | 21 (91%) | 31 (100%) | 31 (100%) | 33 (89%) | 34 (81%) |
| University | 1 (4%) | 1 (4%) | 0 (0%) | 0 (0%) | 3 (8%) | 4 (10%) |
| Highest level of professional teacher training completed |  |  |  |  |  |  |
| Certificate in education (P1/P2/P3) |  |  | 4 (13%) | 0 (0%) | 1 (3%) | 1 (2%) |
| Diploma in Education | 0 (0%) | 2 (9%) | 6 (19%) | 0 (0%) | 4 (11%) | 9 (21%) |
| Degree (Bachelor of Education) | 0 (0%) | 1 (4%) | 0 (0%) | 0 (0%) | 1 (3%) | 1 (2%) |
| Certificate in ECDE | 13 (54%) | 12 (52%) | 9 (29%) | 13 (42%) | 5 (14%) | 8 (19%) |
| Diploma in ECDE | 10 (42%) | 8 (35%) | 12 (39%) | 18 (58%) | 24 (65%) | 20 (48%) |
| Degree in ECDE | 1 (4%) | 0 (0%) | 0 (0%) | 0 (0%) | 2 (5%) | 3 (7%) |
| Employer |  |  |  |  |  |  |
| County | 16 (67%) | 15 (65%) | 31 (100%) | 31 (100%) | 35 (95%) | 39 (93%) |
| BOM | 8 (33%) | 6 (26%) |  |  | 1 (3%) | 2 (5%) |
| Other (NGO/church) | 0 (0%) | 2 (9%) |  |  | 1 (3%) | 1 (2%) |
| Duration of teaching in the current school |  |  |  |  |  |  |
| Less than a year | 3 (13%) | 0 (0%) | 0 (0%) | 1 (3%) | 11 (30%) | 2 (5%) |
| 1 - 5 years | 1 (4%) | 6 (26%) | 11 (35%) | 8 (26%) | 23 (62%) | 33 (79%) |
| More than five years | 20 (83%) | 17 (74%) | 20 (65%) | 22 (71%) | 3 (8%) | 7 (17%) |
| Duration of in teaching profession |  |  |  |  |  |  |
| 1 - 5 years | 0 (0%) | 0 (0%) | 0 (0%) | 1 (3%) | 3 (8%) | 1 (2%) |
| More than 5 years | 24 (100%) | 23 (100%) | 31 (100%) | 30 (97%) | 34 (92%) | 41 (98%) |

### Teachers’ perceptions of the benefits of play in promoting children’s early learning and development skills

Interviewed teachers perceived that play promotes cognitive development and provides opportunities for children to develop new ideas, discover concepts, manipulate games, and retain concepts.

> *"…when you teach through play, children will understand easily, and you will not struggle to teach. Because now, when you want to teach something and introduce it as a story, they will understand and retain it.” **FGD with teachers.***

Generally, most teachers reported that children learn best, understand easily, and become more engaged through play activities such as dramatization and story-telling. These activities enhance the acquisition of new vocabulary.

> *"Learners do understand better when they play and learn simultaneously. For example, when the children are to learn about money and they role-play buying and selling, they will be excited and at the same time learn new vocabularies like money, coins, and notes. When they are playing, the children become active." FGD with teachers.*

Teachers also mentioned that playful activities such as sports promote the development of children’s fine and motor skills, cognitive, and imagination skills. Teachers believe learning through play involving the use of realia, makes children enjoy experiences and promotes mastery of new concepts.

> *“It makes learning and the concept the teacher is teaching real. When they use the materials, they understand the concept and won’t forget it. They develop their imagination by sorting into groups, developing cognitive, small, and large muscle skills through play.” FGD with teachers.*

### Parents/caregivers’ perceptions of the benefits of play in promoting children’s early learning and development skills

As shown in **Figure 1**, over 75% of caregivers/parents from the three counties reported that play promotes social skills. The majority (over 72%) perceived play as key in promoting learning and development. According to some caregivers (**Figure 1**), play was perceived as a contributor to promoting language and cognitive development in young children.

**Figure 1.**
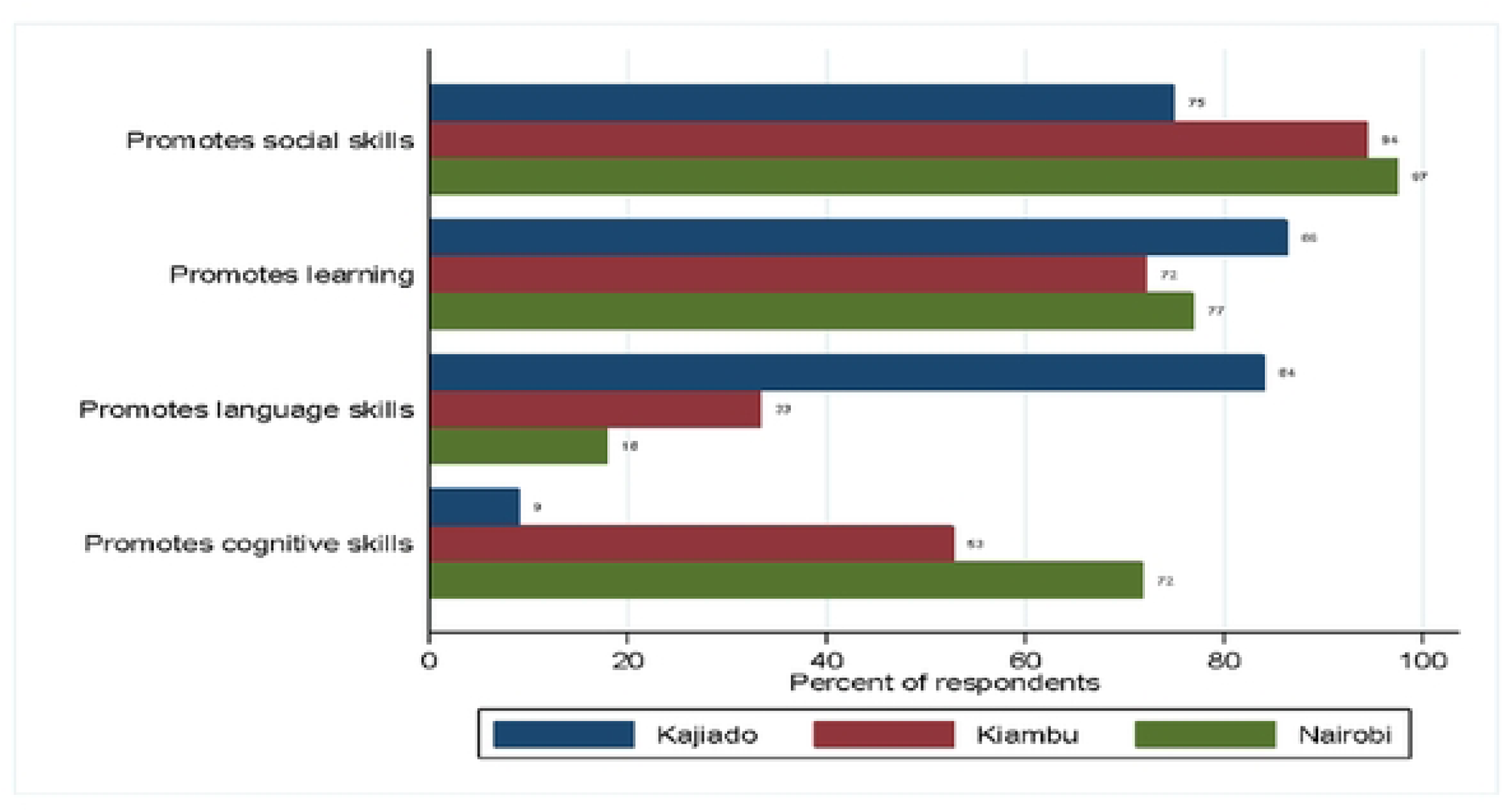
Caregivers’ perceptions of the benefits of play in children

Parents believed that while playing, children learn key concepts such as number work that could later be translated into active learning and practical experiences.

> *"It is through play that she learned. While they play, other children count, and she imitates. Later on, I would help her understand, for instance, the meaning of 5 as a number.” FGD with mothers.*

Similarly, male caregivers perceived that engaging with their children through play was a way of bonding and experiencing fun. They added that playing with children, particularly using devices such as mobile phones, enables children to become relaxed and promotes father-child bonding.

> *“Playing with children is a way of bonding; that’s how we bond, and they can express themselves. When the child is out to play, the muscles become stronger. When the child sees other children happy and playing, the child also becomes happy.” FGD with fathers.*

### Perceived roles of teachers in the implementation of play-based learning approaches

Teachers reported that their roles in promoting LtP included guiding children during role-play activities and linking play with learning activity areas at school.

> *"For example, you can be teaching about jumping; they’re jumping following instructions, and you tell them to jump as they count from one up to ten. So, you have linked mathematics and the outdoors.” FGD with teachers.*

The parents recognized teachers’ perceived roles in play-based learning as they (teachers) ought to have taught the children how to make play materials from locally available resources and ensured children played in a clean and safe environment.

> *"It is important that teachers train pupils to make play materials from locally available materials, such as molding clay. Ensuring that children play in a clean and safe environment and guiding children in all aspects.” FGD with mothers.*

Interviewed fathers suggested that teachers could promote learning through play through storytelling (Bible and other stories), where children would be allowed to ask questions, learn certain concepts.

> *"When narrating stories to them, they are normally not bored; they are very attentive, and, in the process, they do ask questions, and through those questions, I’m able to teach them and reach them. I think this is one way a teacher can promote learning for the children." FGD with fathers.*

### Perceived roles of parents in the implementation of play-based learning approaches

Our findings for the quantitative results revealed that the provision of play materials, play opportunities, and playing with the children were the main areas parents perceived as key contributors to the implementation of play-based learning (**Figure 2).**

**Figure 2.**
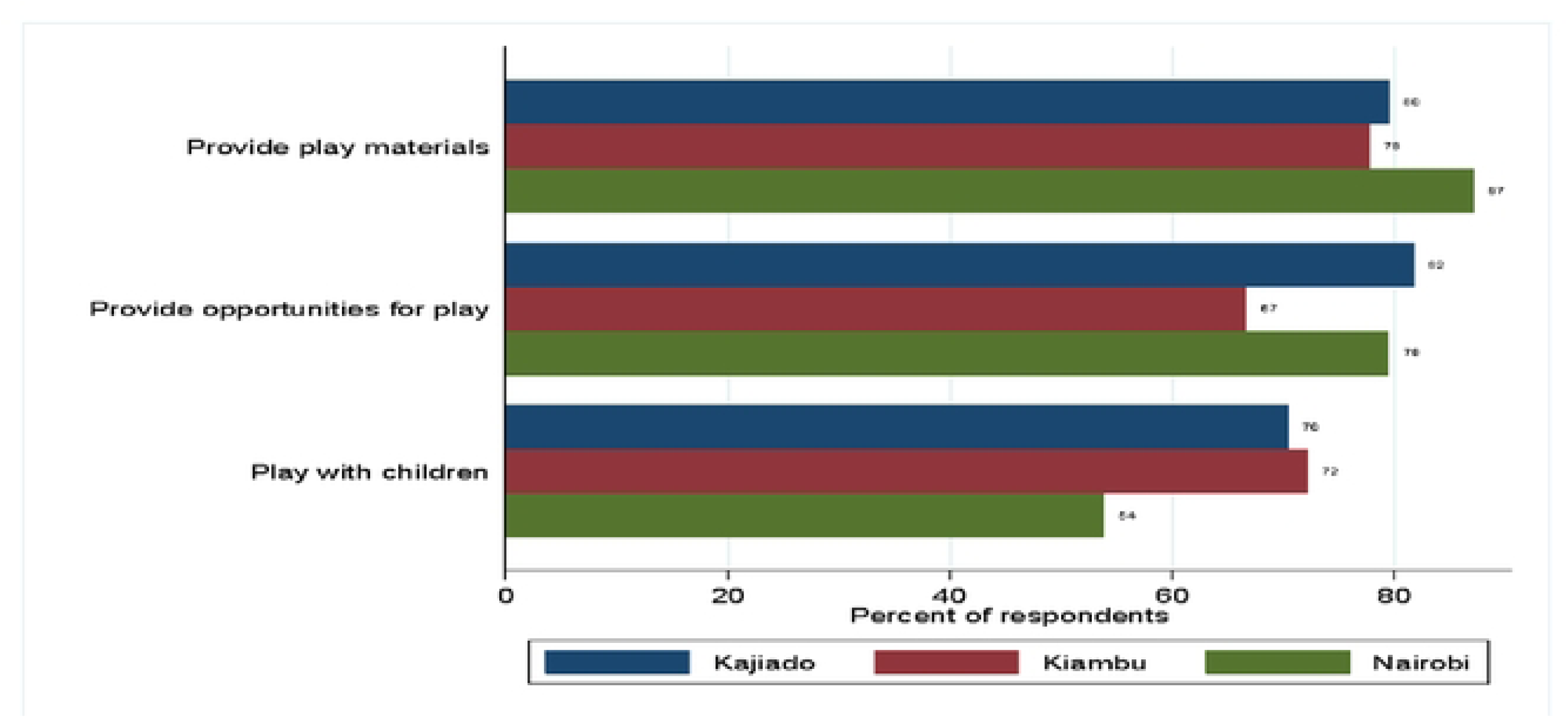
Roles of parents in promoting play-based learning

Most mothers reported that their roles in LtP are to ensure that children are safe and guided during play activities. The mothers also reported that they get involved in their children’s playful activities and are encouraged to pursue their talents even as they provide materials.

> *"I have to be there to make sure they are safe as they play, so as a parent, I have to monitor what they are doing. Encourage and support the child in the games he or she likes because those games alone can take your child to greater heights. I also ensure that they have enough materials to play with.” **FGD with mothers***

## Discussion

In this paper, we sought to explore the caregivers’/parents’ and teachers’ perceptions and roles of play in promoting early learning and development in young children. We wanted to provide insights about the relevant playful learning experiences and how the beliefs and roles of teachers and parents can crystalize play based approach into a meaningful strategy to enhance learning and development in children. Our primary findings revealed that parents and teachers perceived play as an important aspect of children’s lives. They believed that play-based learning contributes to children’s development of fine and motor skills, cognitive, and social skills. Our study results show that parents’ and teachers’ availability and involvement during playful learning were considered critical in promoting children’s learning and development of specific skills and mastery of learning concepts. The parents believed that teachers were required to provide opportunities for learners to use locally available play materials to construct playful experiences, learn from them, and explore.

### Teachers’ perceptions of the benefits of play

Results indicated that teachers perceived play as crucial in promoting learning and development in children. Playful learning experiences were believed to promote cognitive skills, fine motor skills, and acquisition and mastery of new concepts. Further, LtP was considered an important interactive instructional strategy that would enhance the development of new and emergent vocabularies.

Our findings revealed that there was a generally positive perception and agreement from teachers that children loved to play and they would never get bored playing and learning. Our study findings mirror that of a mixed-method approach research conducted in Palestinian schools, whereby interviewed teachers recognized the value of play-based learning and were highly motivated to incorporate it as an instructional pedagogy (Khalil et al., 2022). Teachers from ECE centers in Zambia expressed a positive perspective on play-based learning asserting that play pedagogies benefit children holistically. However, the implementation of play-based learning in Zambian ECE centers varied. For instance, some teachers engaged children in play activities only during scheduled time for play while other teachers integrated play into the delivery of lessons (Lungu & Matafwali, 2020). These findings add new knowledge to our research including the motivations behind teachers’ perception of play-based learning, training skills, and the significance of varied contexts and factors that contribute to the use of play pedagogies.

### Parents/caregivers’ perceptions of the benefits of play

Parents are the first teachers of children. Their perceptions of the use of play-based learning impact on the way children learn and develop. Our research findings showed that parents recognized play as a contributor to children’s learning and development across social, language, and cognitive skills. Implementation of play-based learning was considered more effective through joint parental and child engagement. This allows children to relax and view play as a thrilling activity. The results concur with a study conducted among parents from Southwestern City in the U.S. which asserted that play-based learning comprises positive and negative perspectives. Parents view play as a contributor to social interaction, and fun, and presents opportunities for creativity and exploration in children (Ducusin & Dy, 2016).

Negative aspects of play discovered by parents included injuries, imitation of bad behaviors, and engaging in too much digital or screenplay (Id et al., 2024). Parents from a study conducted in Ethiopian preschools viewed play as a contributor to their children’s academic and cognitive skills (Metaferia et al., 2021). Further, children’s increased frequency of having breakfast was perceived as a predictor of enhancing play-based learning (Metaferia et al., 2021). Parents also perceived socio-cultural variations and socio-economic factors as contributors to the use of play in promoting executive function in children. These findings provide additional knowledge to this paper, particularly evidence that play has multiple socio and cultural predictors at the family level, which could be a potential area for researchers to explore in future studies.

### Perceived roles of teachers in the implementation of play-based learning approaches

Our results revealed that teachers plan play spaces and contribute to the development and provision of play materials. Teachers also guide and link playful activities with learning activity areas in school. The results concur with a study conducted in Thailand preschool centers which found teachers as planners, communicators, reflectors, and observers during children’s playtime (Jannah & Asmara, 2023). In sub-Saharan Africa, specifically Kenya, teachers in preschools did not adequately integrate play as an instructional pedagogy, inhibiting children from exploring playful learning (Murundu et al., 2014). This environment implies that learners would mostly miss out on the essential executive skills requisite for school readiness and life learning.

LtP perception is reinforced by Lunga et al. (2022), who state that teachers view play as a vital tool for creating a less stressful learning environment and enhancing child development. However, this could differ significantly, particularly between structured and unstructured play. Moreover, teachers’ lack of adequate knowledge, skills, and competencies to integrate play as a key strategy to attain the later skills warrants more research into this area to explore the implementation of play-based learning in preschools. Research also shows that some teachers advocate for guided play activities that align with educational goals, while others favor the open-ended exploration offered by free play (Timmons et al., 2021). Whereas our study findings demonstrated that teachers play an important role in guiding children, the results from the later research add a new body of knowledge to this paper and largely attract more scientific evidence to this topic.

### Perceived roles of parents in the implementation of play-based learning approaches

In a fast-evolving early childhood education, play-based learning is considered critical in promoting children’s development and learning. Our findings in this paper highlight parents’ engagement during play, provision of play materials, and facilitating a safe play environment as important discourse in promoting LtP. Whereas the results of a study conducted in low-income families of African-American Latino families revealed parents’ acknowledgment that play promotes children’s school readiness and social skills, our paper did not establish this (LaForett & Mendez, 2017). Further, the results indicated how culture interacts and can influence the use of play as well as the alignment of play-based pedagogies with the home setting (LaForett & Mendez, 2017). Interestingly, the findings imply that culture may affect the attitudes of parents and potentially influence their contribution to promoting playful learning; a new area that can be investigated in the future.

Whereas our findings indicated that parents get involved in play activities to understand the talents of their children and encourage them, results from a study carried out in South Africa revealed that while parents believed that play was important, often they set up structured learning environments for their children (Bipath et al., 2022). This finding contradicts the expectation of societies towards adults’ roles in facilitating children to explore through play. In a study conducted in Central Kenya, parents and family members were considered enablers in promoting a safe physical play environment at home (Ndani, 2019). Free play activities can be dangerous in the absence of adults; thus sensitization of parents and adult family members is important to prevent injuries and promote learning (Bipath et al., 2022; Ndani, 2019). The results from the later studies add new knowledge on the concept of the roles of parents, but also present opportunities for researchers to explore the parents’ capacity needs in facilitating safe learning through play.

Synthesis of our reviewed literature indicates that barriers to effective implementation of play-based learning are not adequately researched in sub-Saharan Africa, particularly among preschools in Kenya. The lack of standardized guidelines and policies on play-based learning and its integration into early childhood education is a major obstacle (Dowd & Thomsen, 2021). Our findings in this paper assert the emphasis parents and caregivers should place on play and their primary responsibility to guide children and present playful opportunities.

### Conclusions and Implications

In our understanding of the perception and roles of parents and teachers in the implementation of play-based approaches, several themes emerged related to convergent and divergent views on play-based approaches. Teachers asserted that play-based learning was a significant instructional strategy and an aspect that promotes the cognitive development of emergent vocabularies in children. Interestingly, some teachers in sub-Saharan African ECE centers perceive play as crucial through integrating it only during playtime time and other teachers integrated play during the delivery of lessons. Based on our results, we encourage preschool teachers to emphasize parental involvement in play and providing emphasis on the benefits of play at home. In addition, we encourage teachers to seek varied capacity-strengthening opportunities to equip them with knowledge about the use of play as an instructional strategy not only during learning but also through promoting explorative safe play activities in children. Findings in this paper support early childhood practitioners’ beliefs that play is energetic, constructivism, and an integral part of children’s development and learning. Since parents are considered first teachers, preschool teachers may identify material development days on play-based learning that involve parents, children, and teachers. This strategy would promote a positive perception and view of the community about the value of play in children and later support teachers in the development of culturally relevant play materials.

Further research may be conducted in similar low-and-middle-income countries to examine teachers’ perception of the use of play as an instructional approach at the teacher training college, father versus mother involvement in play-based learning, young children out of school and older children out of school and observe the role of family members and relatives on play-based learning activities. This research might provide insight to the Ministry of Education (MoE) of Kenya to contextualize and adapt the teacher training curriculum to be used at the training colleges and promote collaboration between parents and teachers on the implementation of play-based learning.

## Acknowledgements

This research was supported by the following authors Paul Otwate who led the conceptualization of this paper and overall leadership in drafting the manuscript. We would like to sincerely appreciate Patricia Kitsao-Wekulo, Margaret Nampijja, and Silas Onyango for providing intellectual input to this paper. Also, we wish to thank Nelson Langat, Silas Ooko, and Brian Odhiambo for leading in writing methodology section. Linda Oloo, Jesse Mabongah, and Erick Makhapila supported literature review.

